## Supplemental Figures for "A bacterial genome and culture collection of gut microbial in weanling piglet"

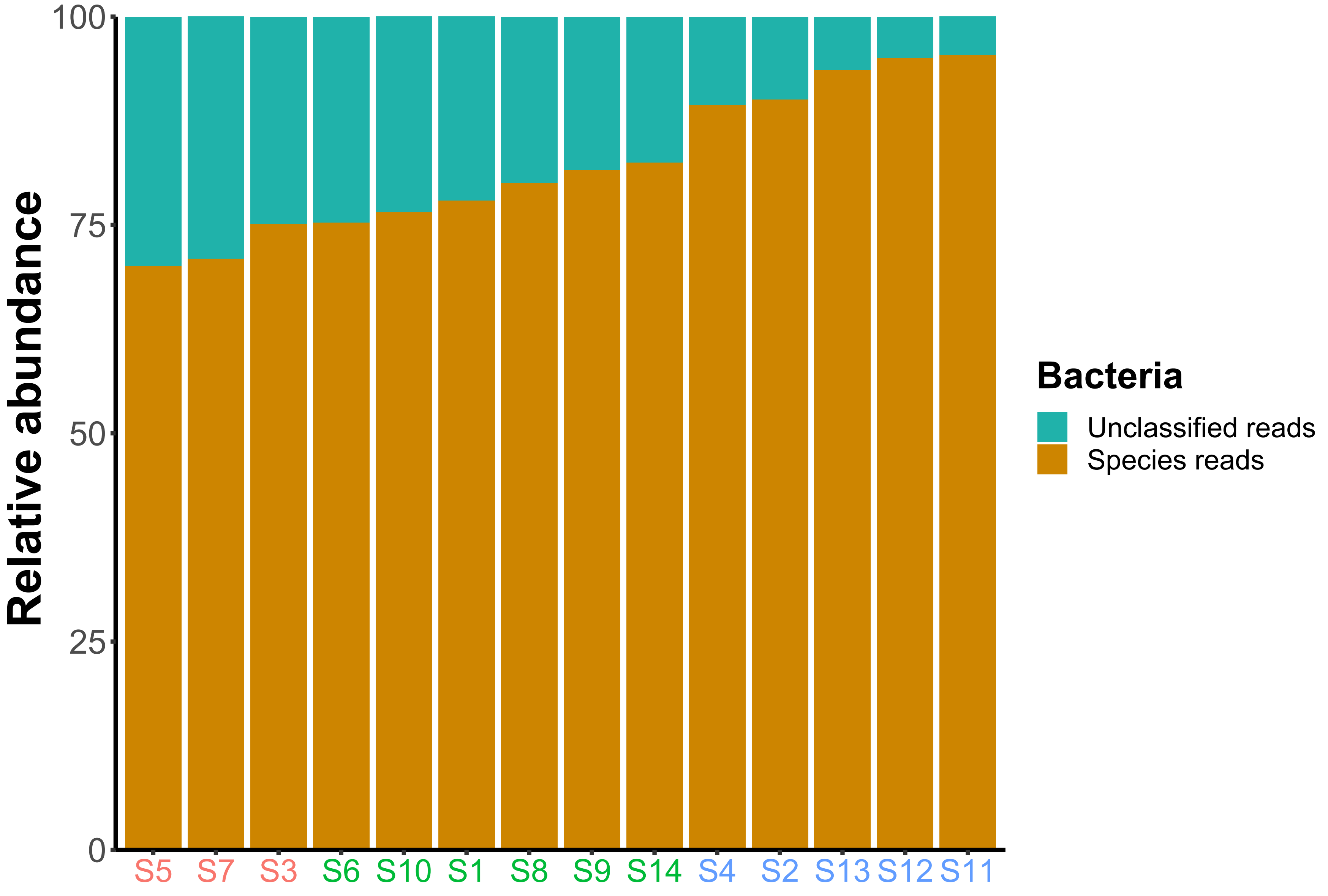


**Supplementary Figure 1**

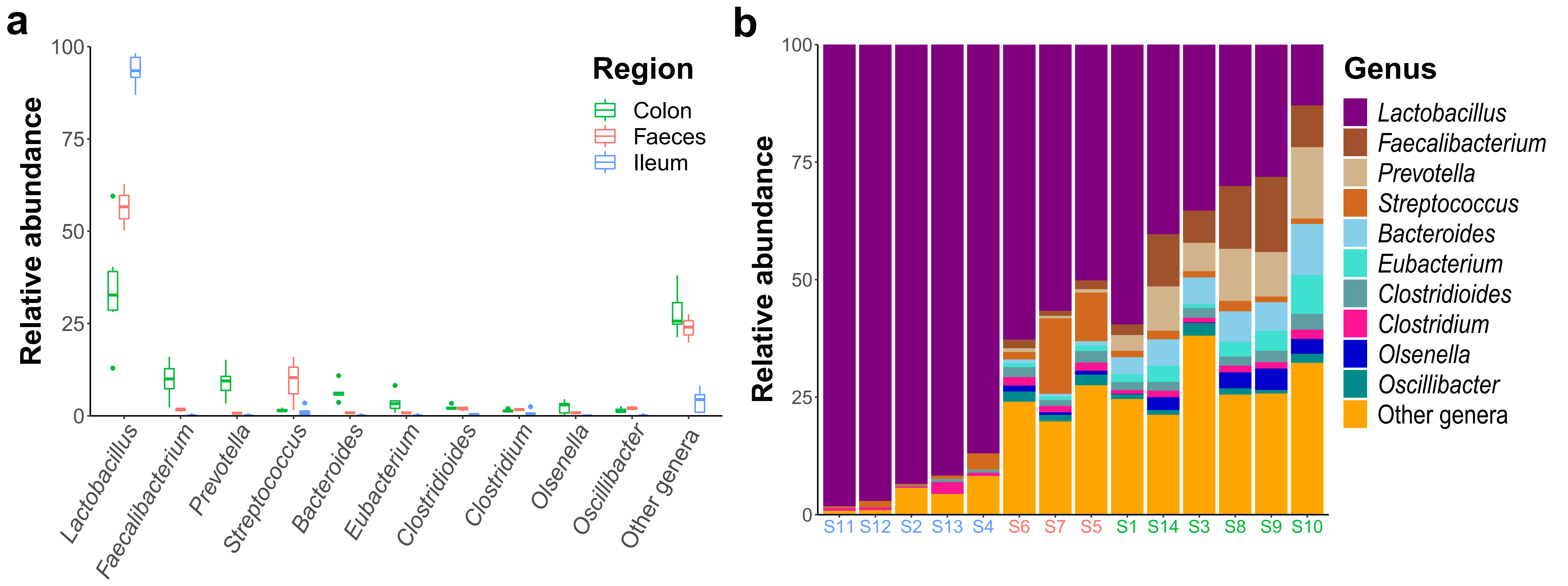


**Supplementary Figure 2**

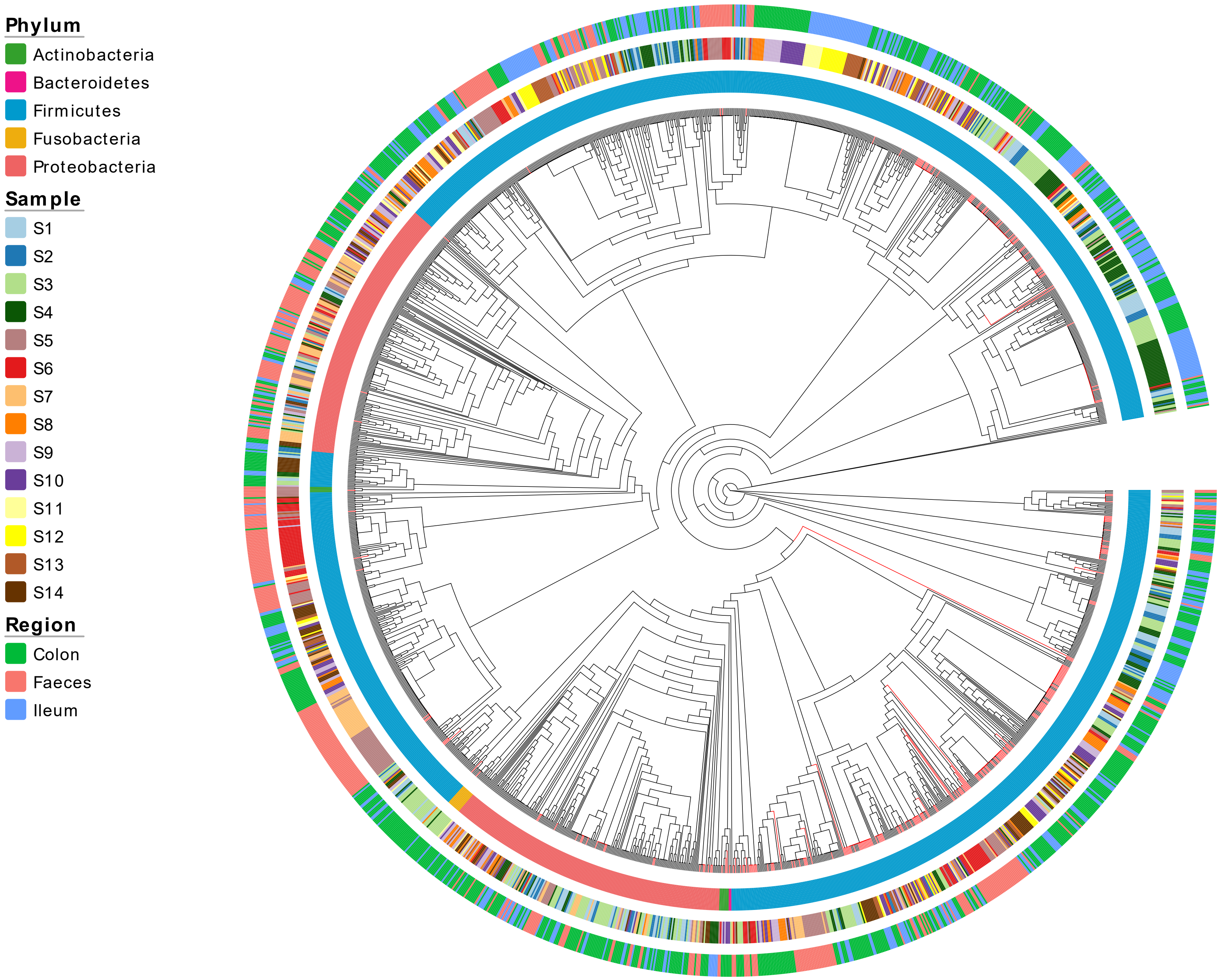


**Supplementary Figure 3**

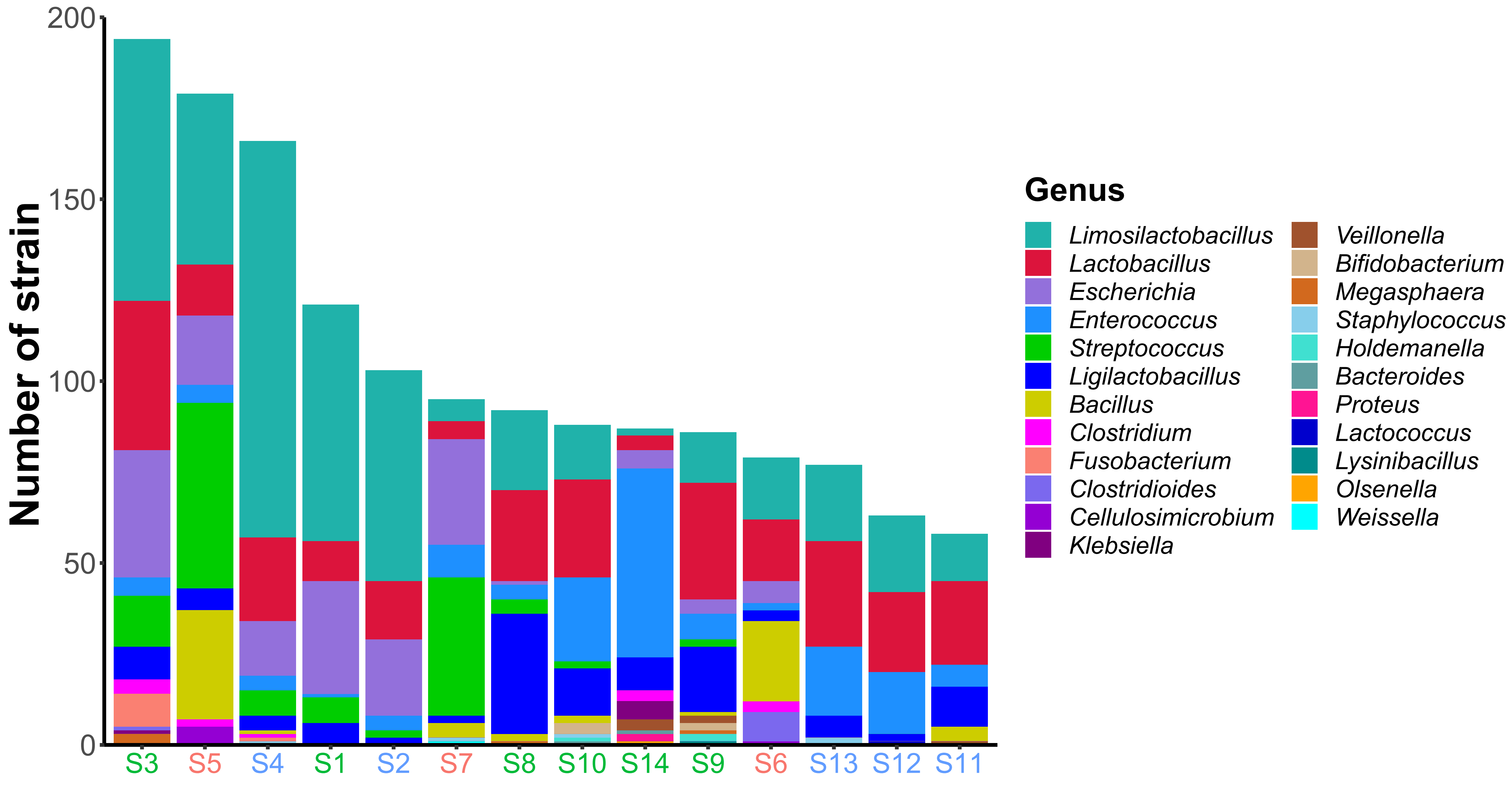


**Supplementary Figure 4**

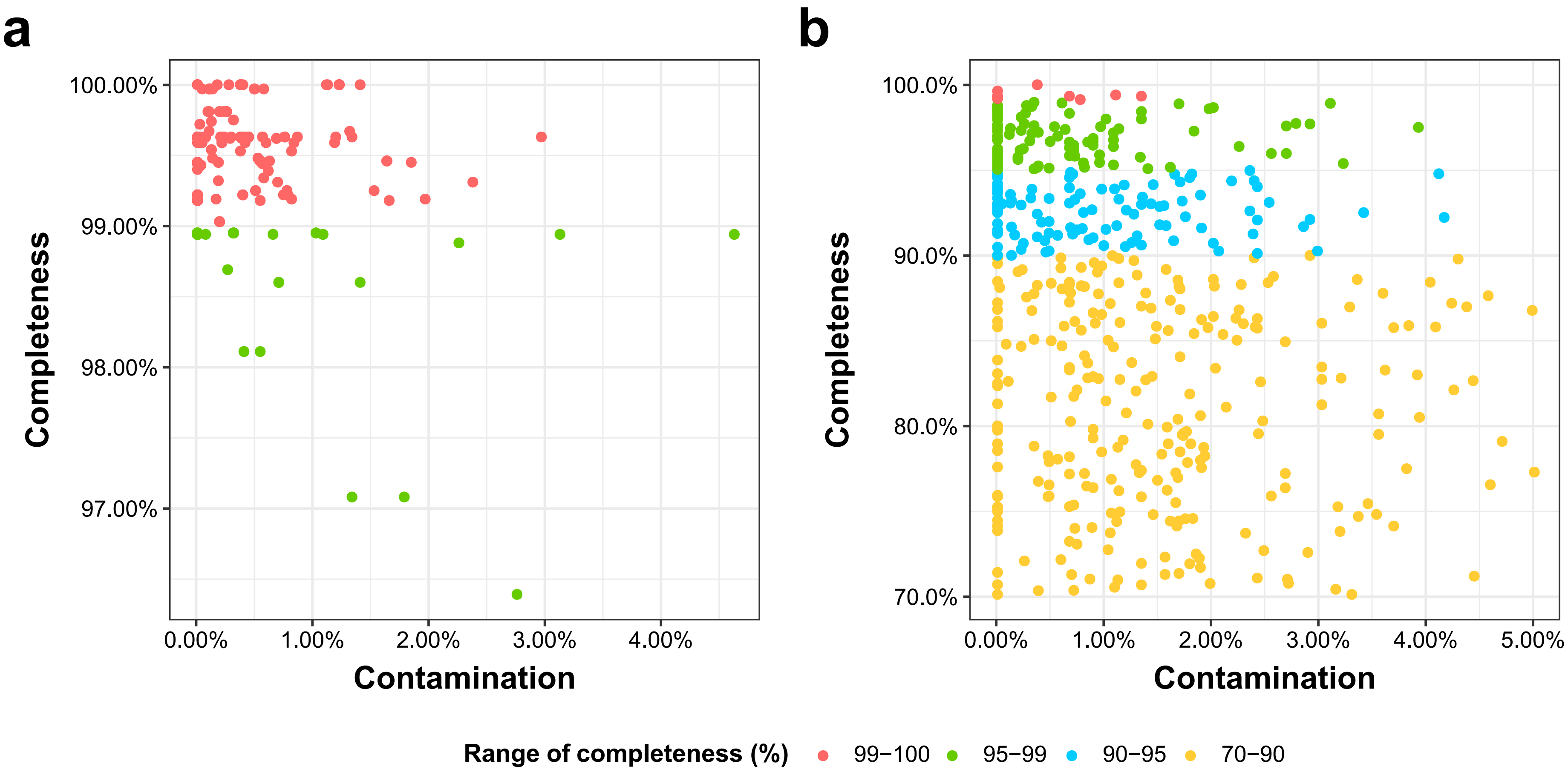


**Supplementary Figure 5**

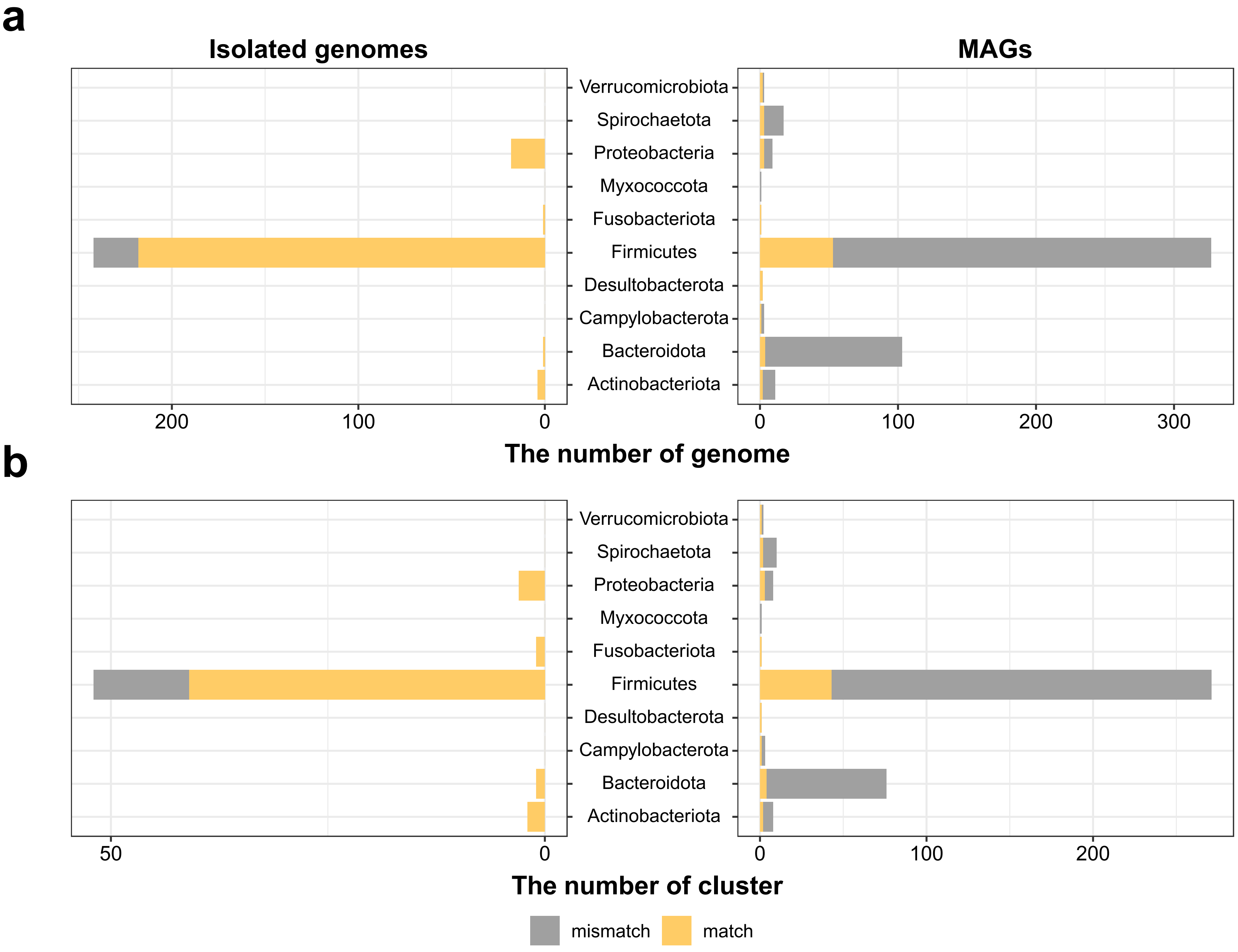


**Supplementary Figure 6**

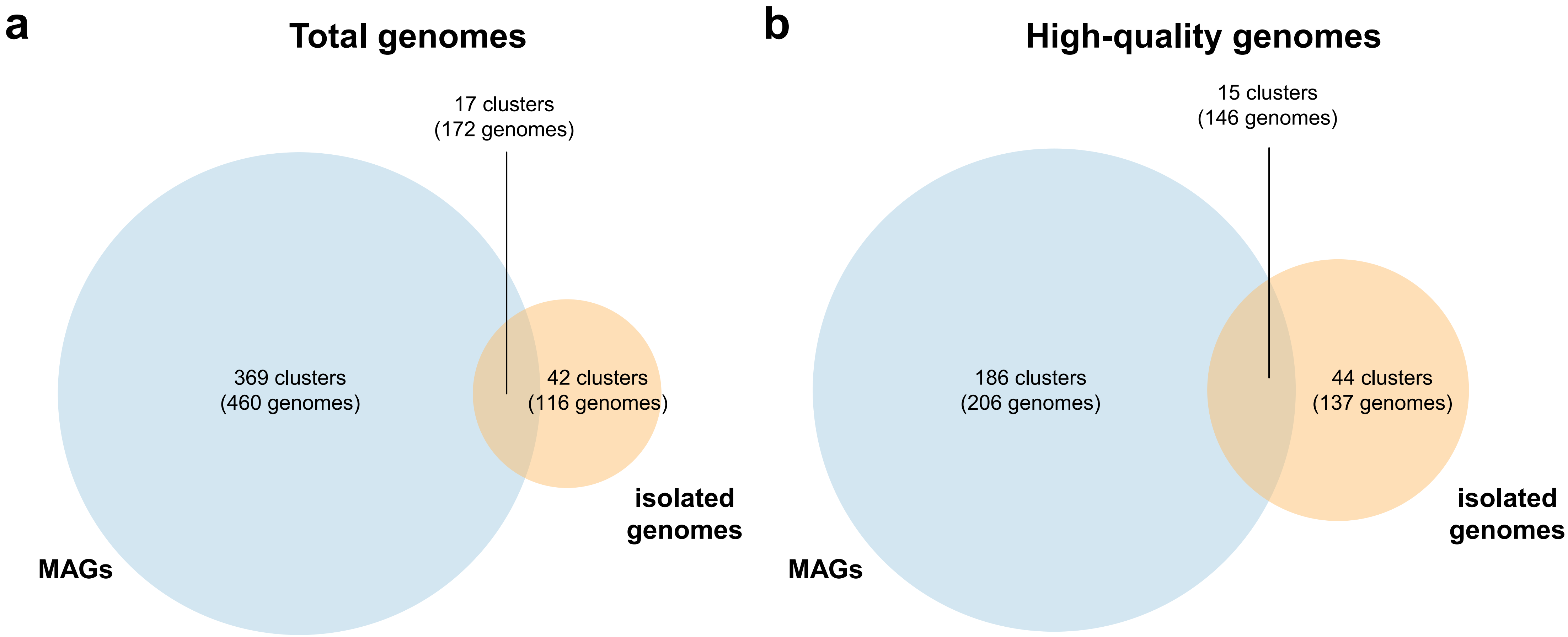


**Supplementary Figure 7**

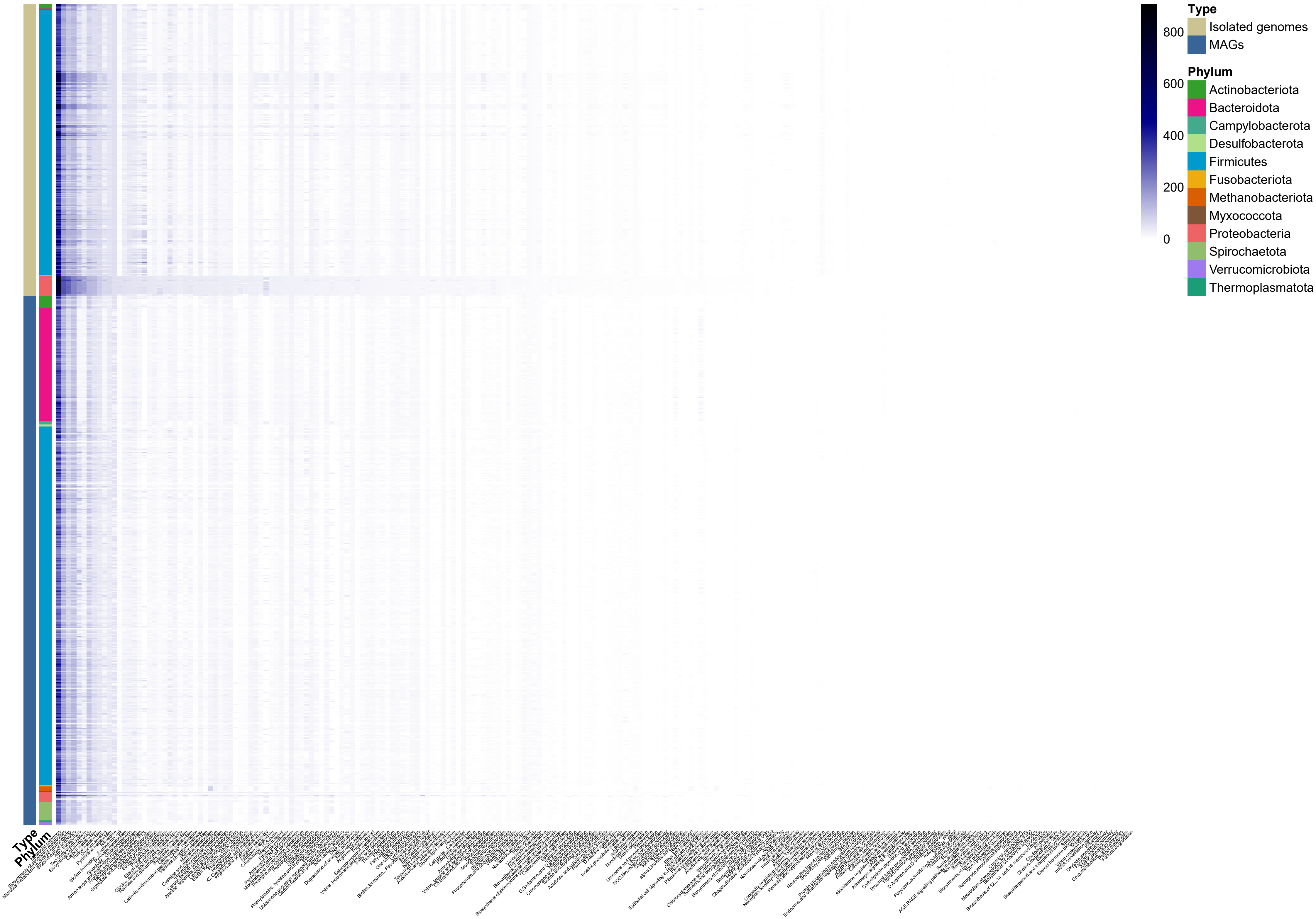


**Supplementary Figure 8**

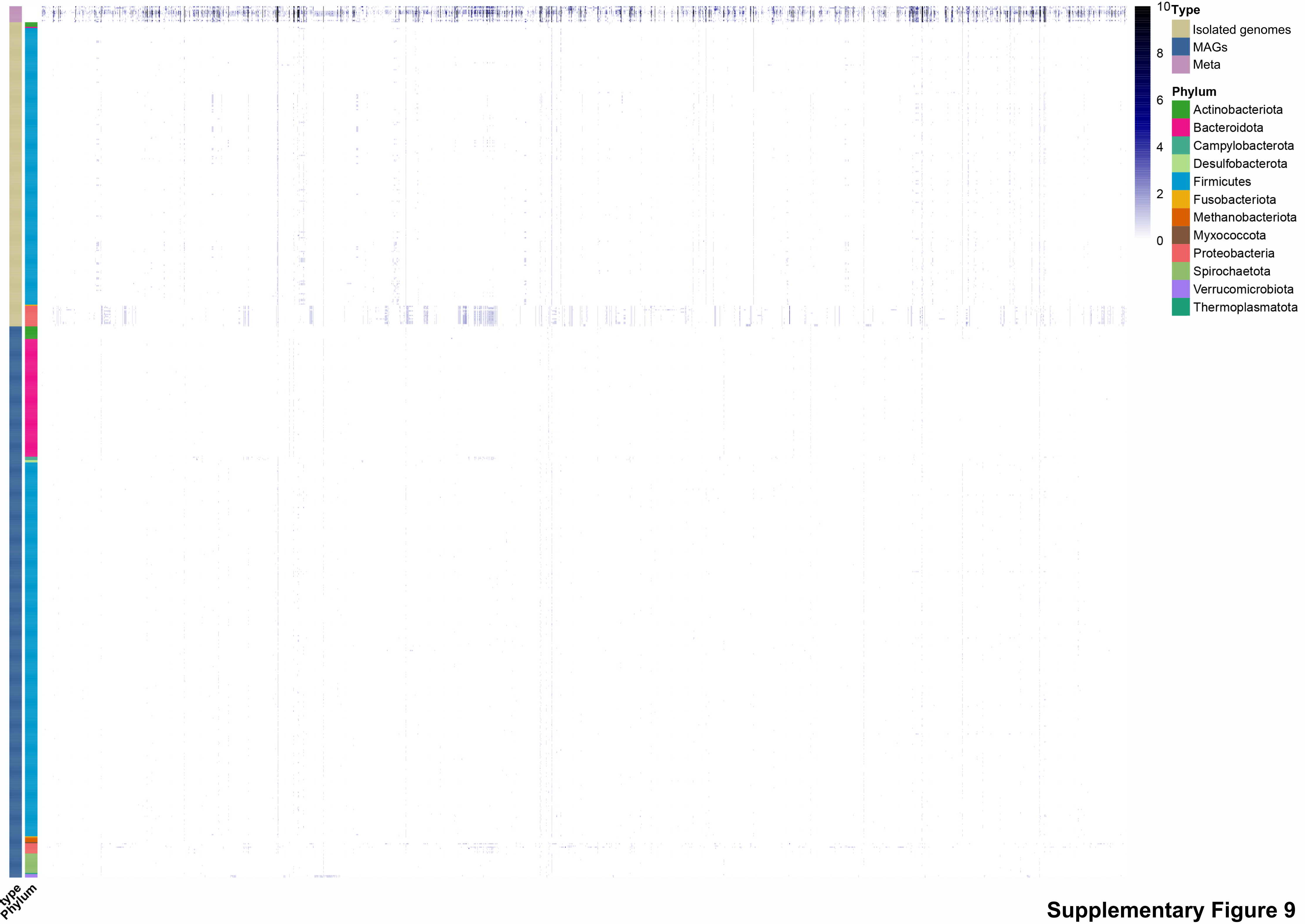


**Supplementary Figure 9**

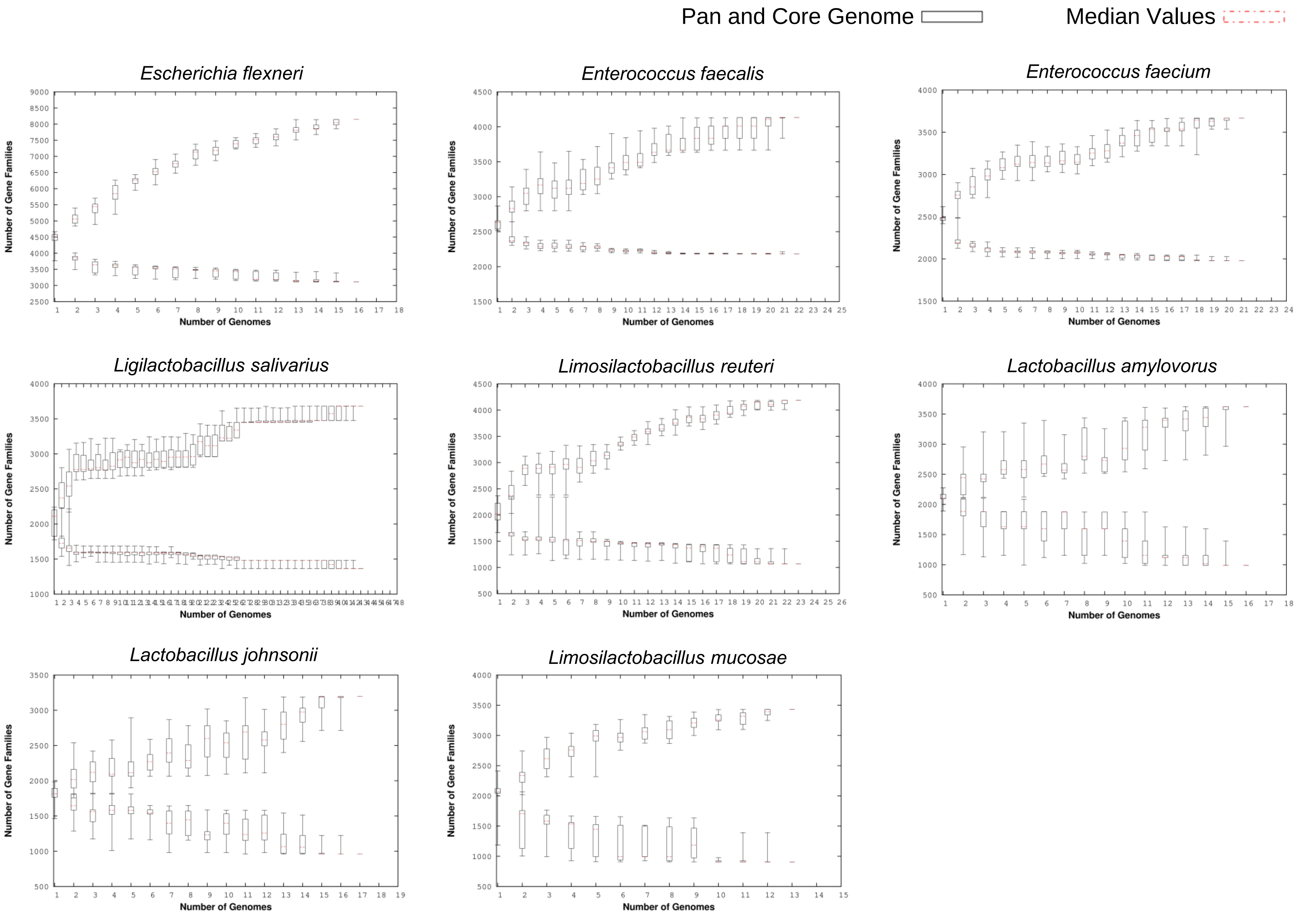


### Supplementary Figure 10

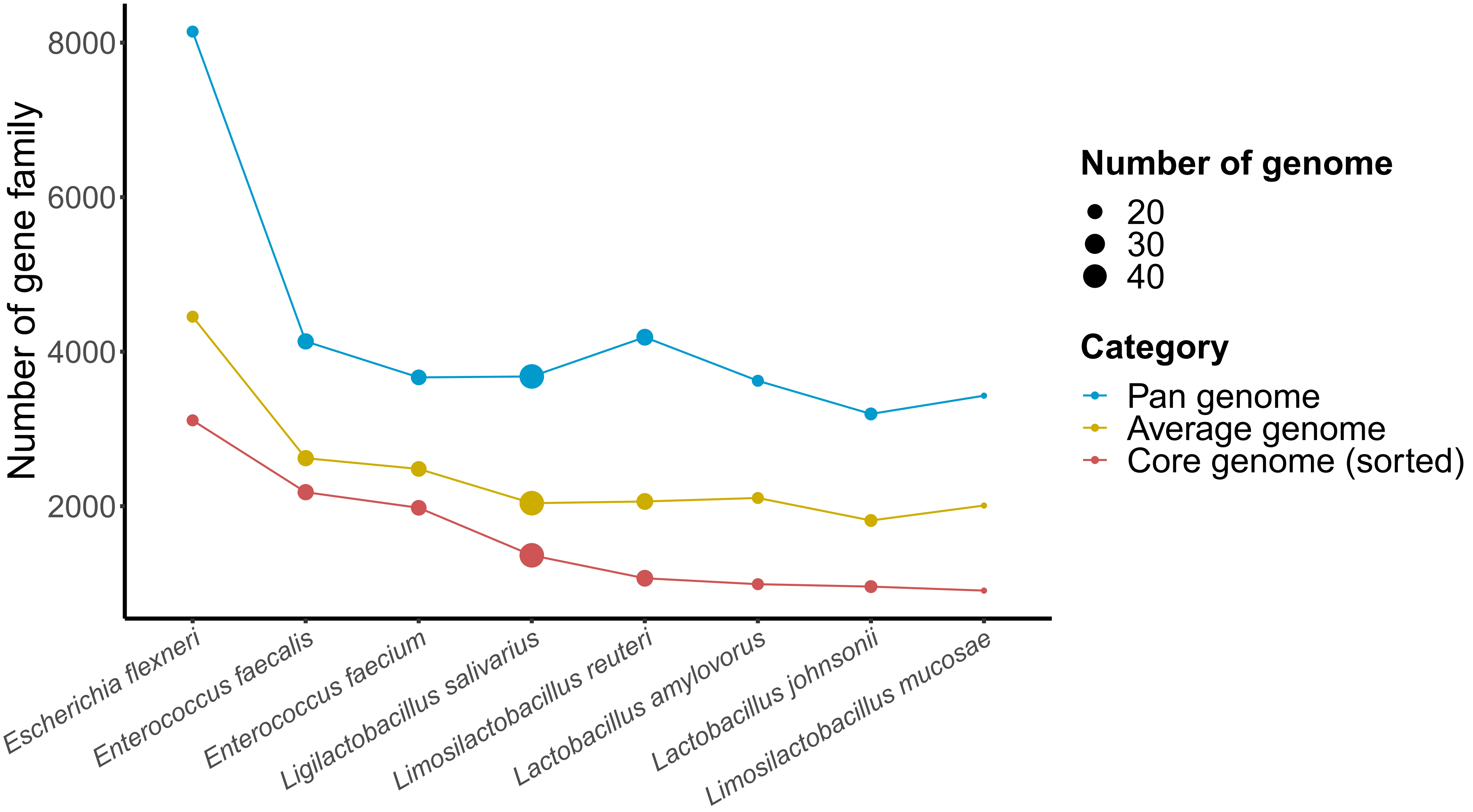


**Supplementary Figure 11**
