## Supplemental Material for "A bacterial genome and culture collection of gut microbial in weanling piglet"

**Supplementary Figure 1. The proportion of reads that were and were not classified at the species level.**

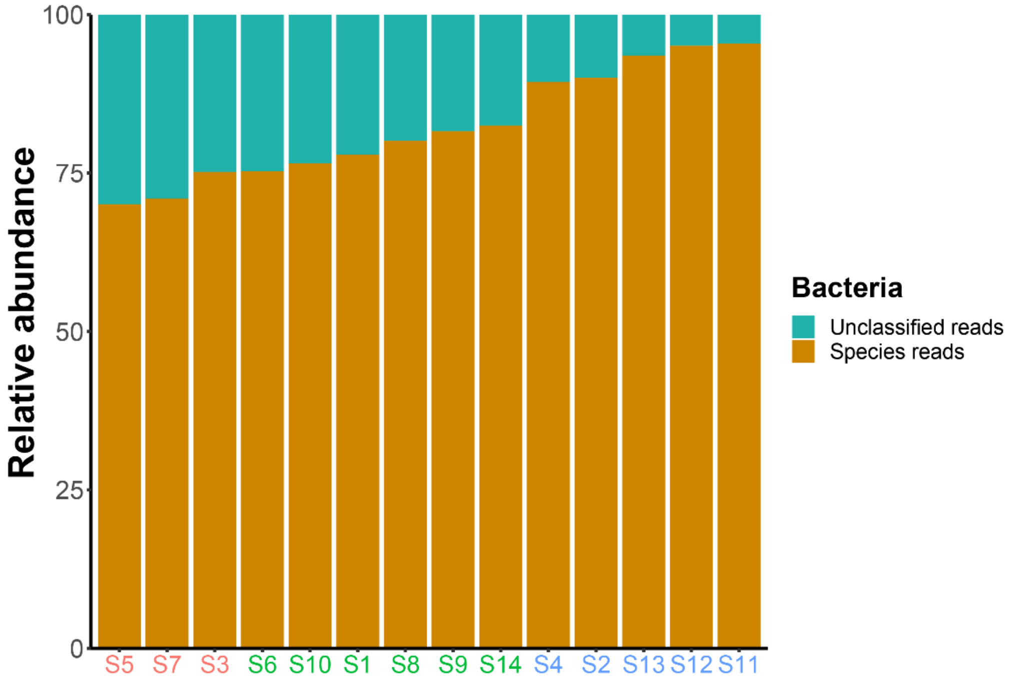


**Supplementary Figure 1**

**Supplementary Figure 2. a-b, The comparison of genus-level proportional abundance in the ileum, colon, and faeces of weanling piglets.**

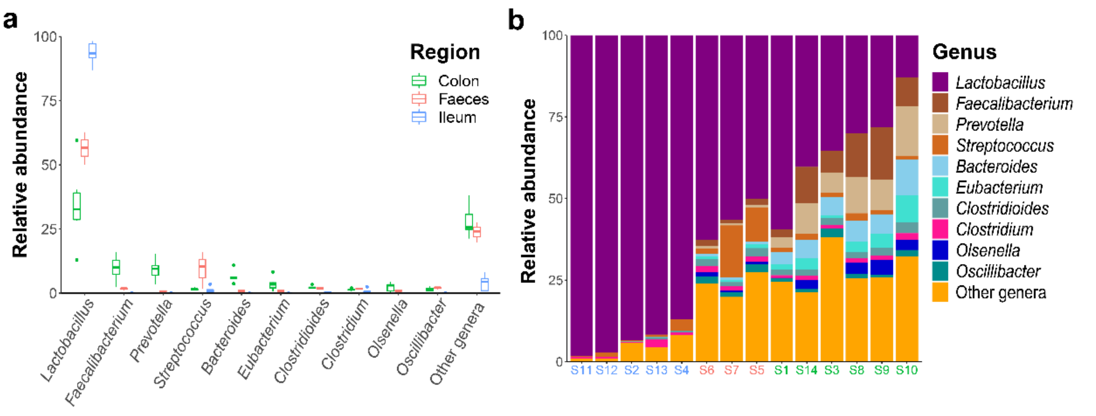


Supplementary Figure 2

**Supplementary Figure 3. The 16S rRNA gene sequence phylogenetic tree of 1,476 strains.** Novel clusters are highlighted by red clades. Phylum, sample, and region are display in the first, second, and third outer layer, respectively.

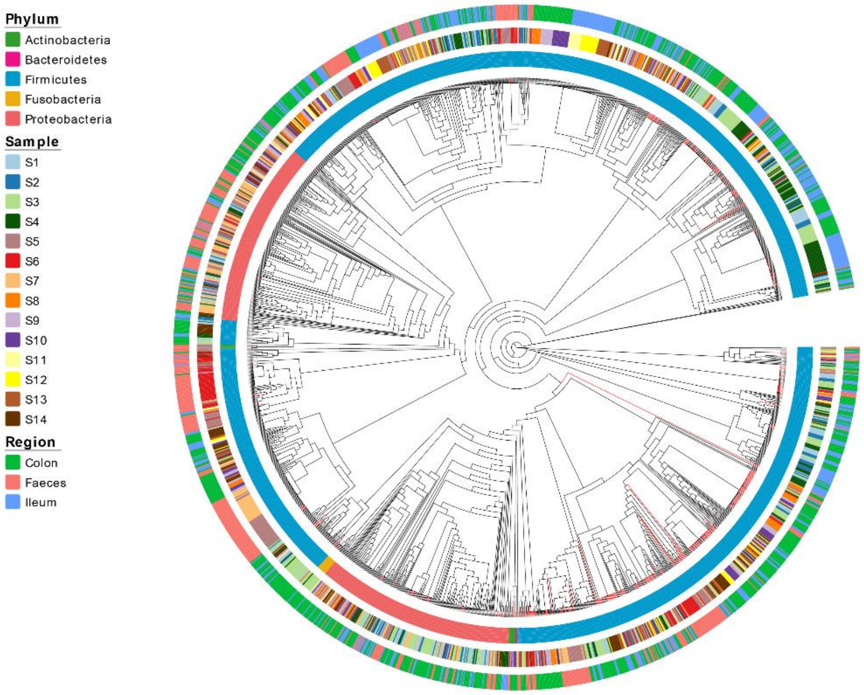


Supplementary Figure 3

**Supplementary Figure 4. The number of cultivated bacterial strains at genus level from 14 samples.**

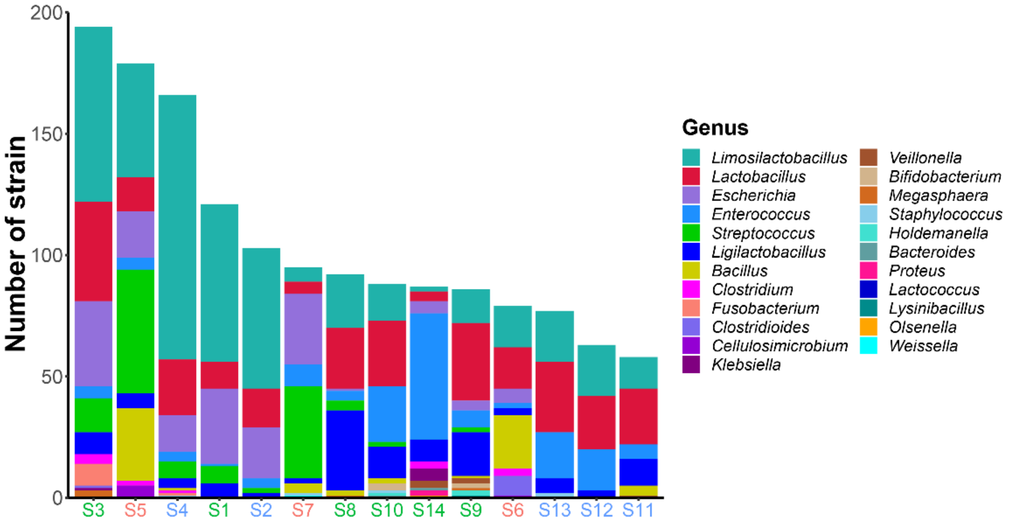


Supplementary Figure 4

**Supplementary Figure 5. Quality assessment of the genomes. a-b,** Completeness and contamination of 266 isolated genomes (a) and 482 MAGs (b), respectively.

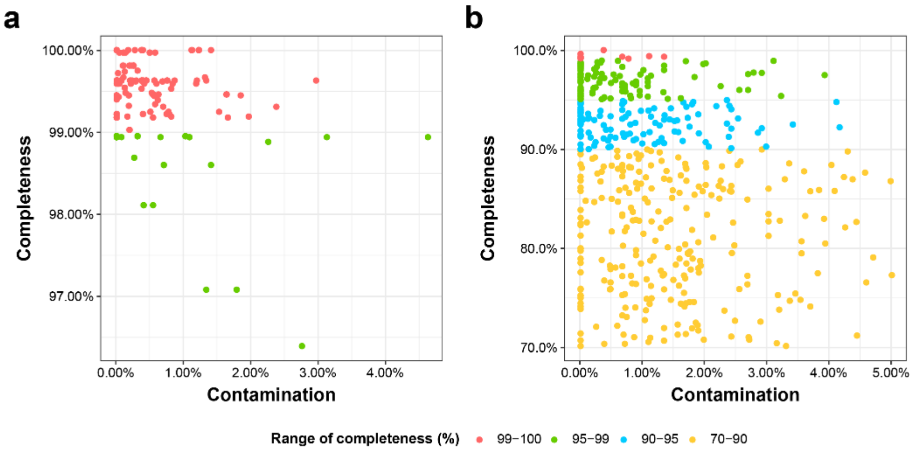


Supplementary Figure 5

**Supplementary Figure 6. The taxonomy of the 743 bacterial genomes. a-b,** The distribution of isolated genomes and MAGs across different phylum. Statistics show the number of genomes (a) and clusters (b), respectively.

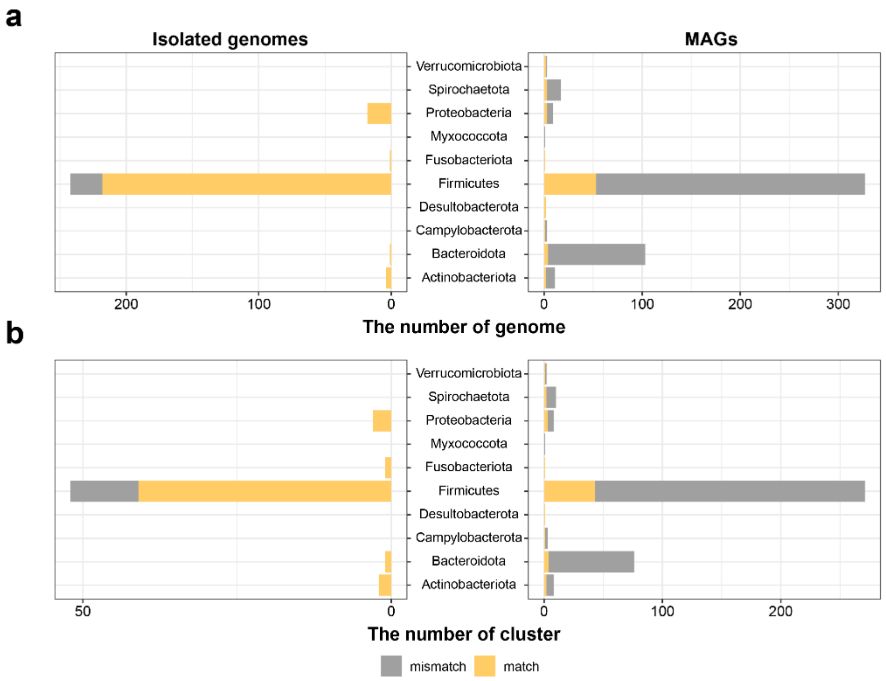


Supplementary Figure 6

**Supplementary Figure 7. Comparison of MAGs and isolated genomes in different quality standards. a,** MAGs with >70% completeness and <5% contamination, while isolated genomes with >90% completeness and <5% contamination. **b,** Both MAGs and isolated genomes are high quality (with >90% completeness and <5% contamination).

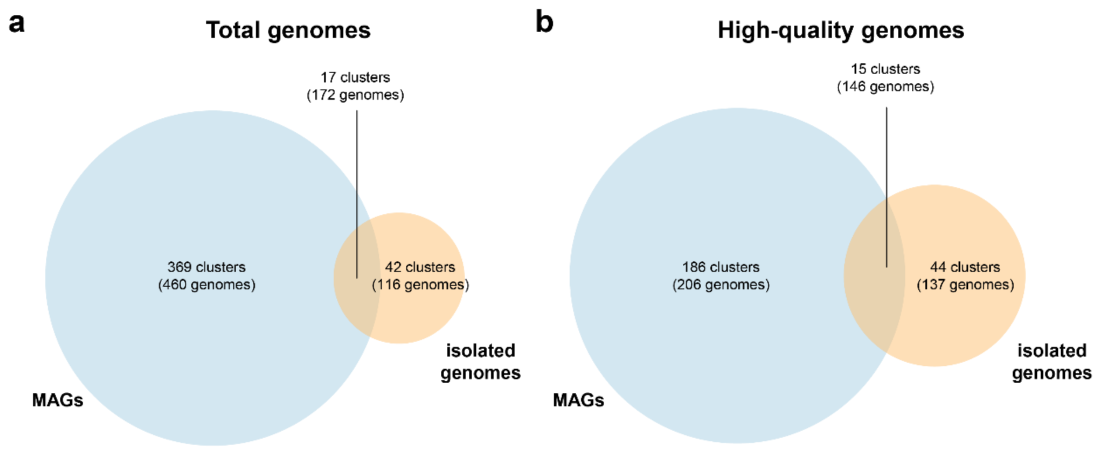


Supplementary Figure 7

**Supplementary Figure 8. The distribution of KEGG pathways in each genome.** The presence and absence of each pathway is indicated by blue and white, respectively.

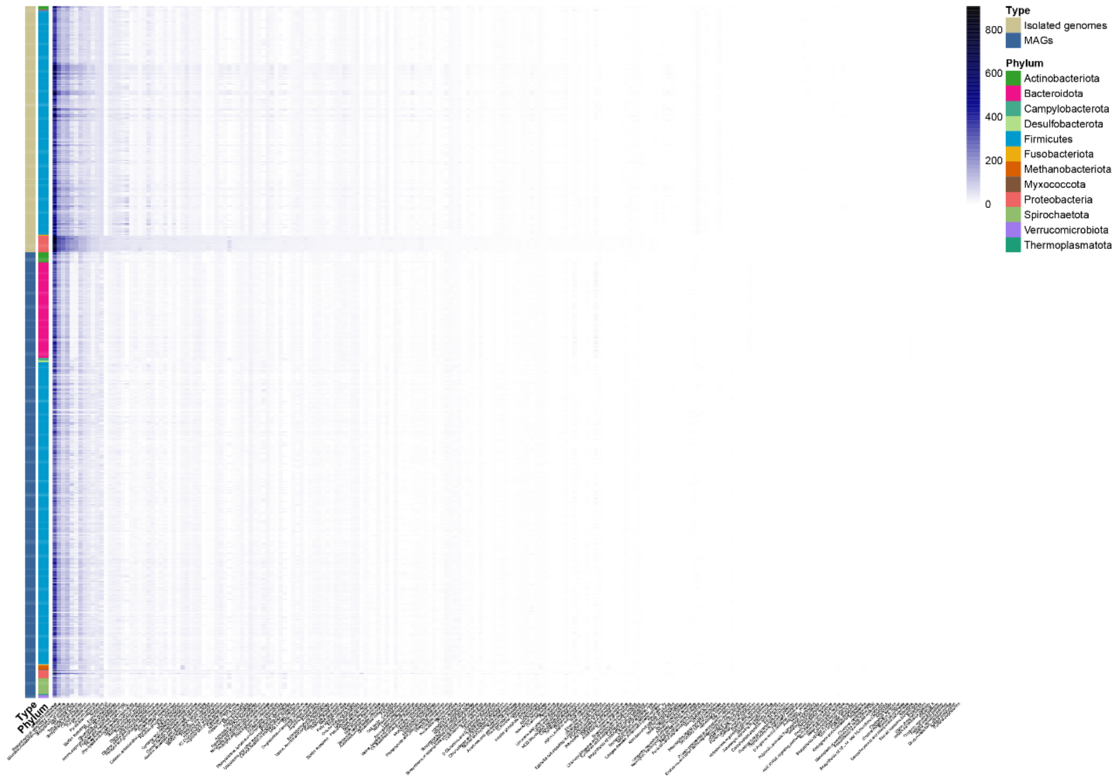


Supplementary Figure 8

**Supplementary Figure 9. The distribution of VFDB in each genome.** The presence and absence of VFDB is indicated by blue and white, respectively.

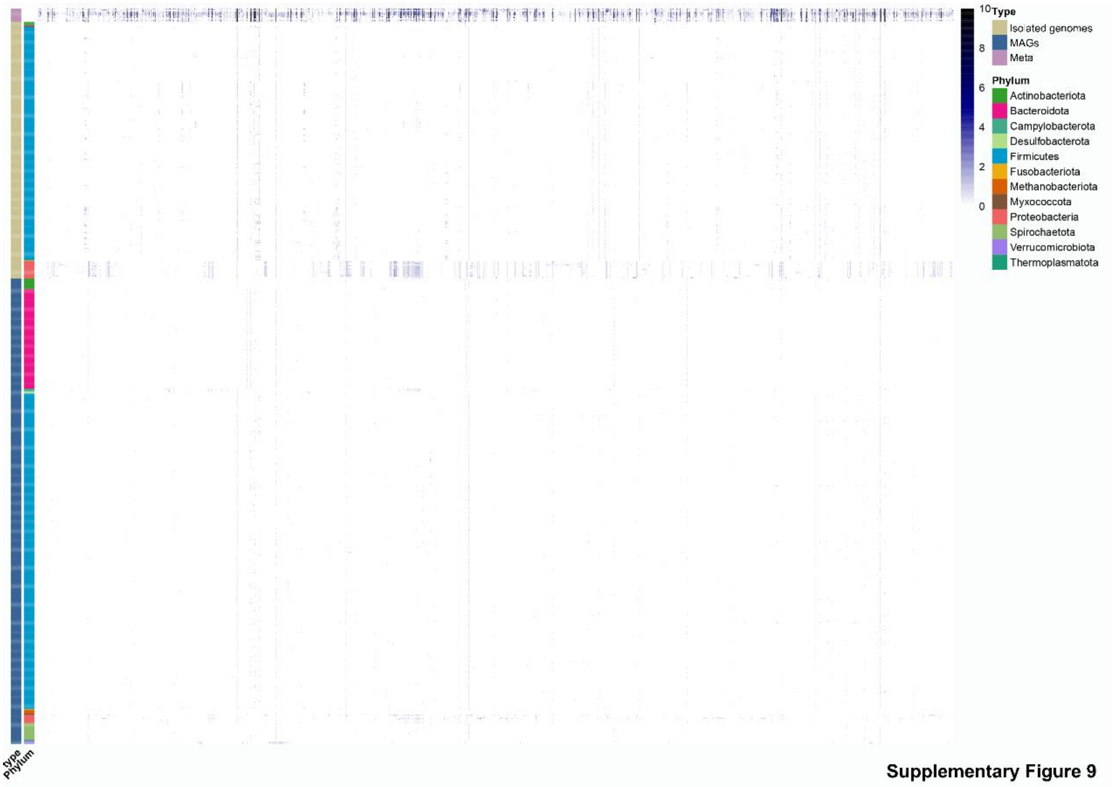

**Supplementary Figure 10. The accumulation curves of the pan and core genome of the 8 representative clusters.** Box plots indicate 25th and 75th percentiles with medians shown as horizontal lines and whiskers set at 10th and 90th percentiles.

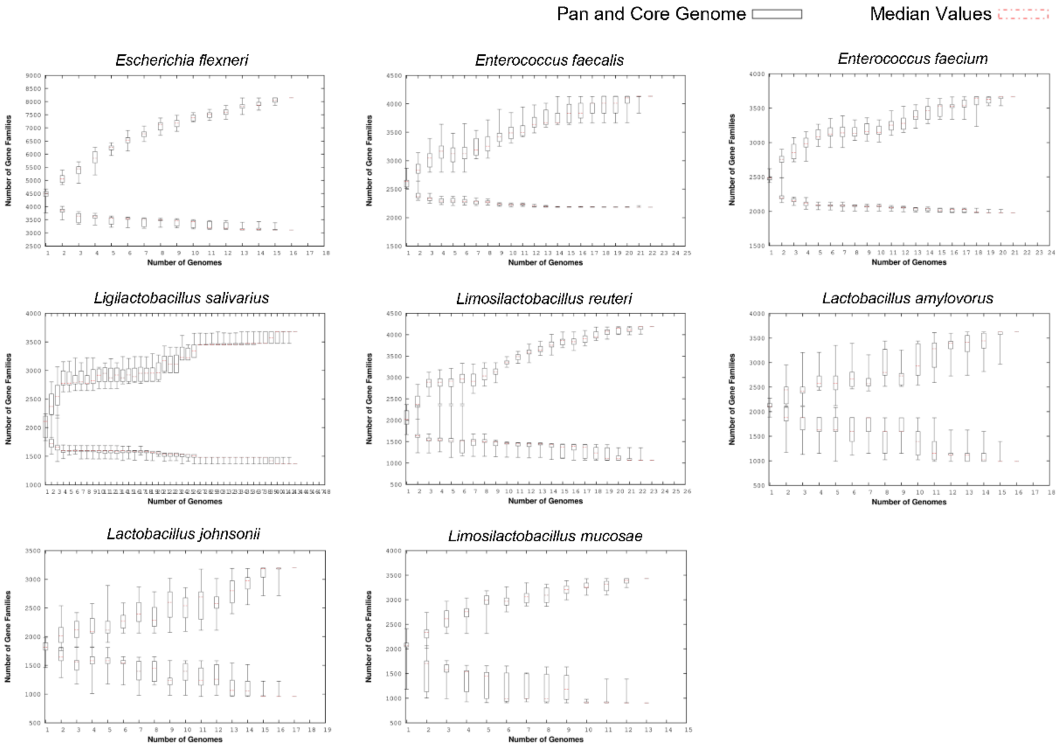


**Supplementary Figure 10**

**Supplementary Figure 11. The gene family numbers of pan genome, core genome and average genome of the 8 representative clusters.**

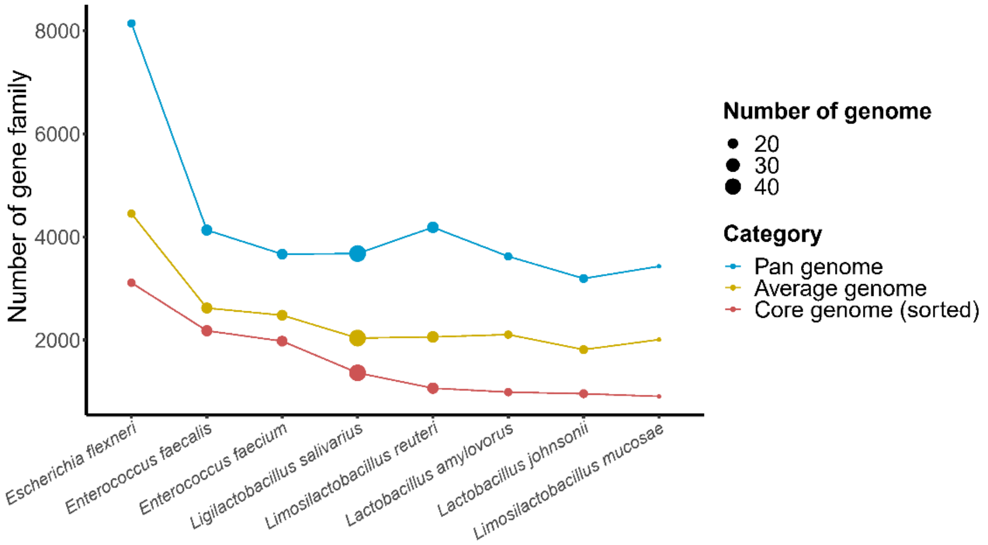


Supplementary Figure 11
