## Supplemental method for "A bacterial genome and culture collection of gut microbial in weanling piglet"

**Supplementary method**

A total of 25 kind of culture media were used for bacteria culture of gut of piglets. The samples were suspended with PBS supplemented with 0.1% cysteine and serially diluted with tenfold in an anaerobic chamber (Bactron Anaerobic Chamber, Bactron IV-2, Shellab, USA). The diluted suspension was then spread on agar plates and incubated under anaerobic condition with gas flow composition of 90% N_2_, 5% CO_2_ and 5% H_2_ at 37 °C.

The culture media used in this study were listed as below:

*5% sheep blood-**MPYG: MPYG+5% sterile defidrinated sheep blood*

*MPYG：MPYG Medium*

*Kana- MPYG: MPYG Medium+ Kanamycin*

*5% sheep blood-BHI: Brain Heart Infusion Medium +5% sterile defidrinated sheep blood*

*BHI：Brain Heart Infusion Medium*

*Kana-BHI：Brain Heart Infusion Medium+ Kanamycin*

*5% sheep blood-DM：DM Medium +5% sterile defidrinated sheep blood*

*DM：DM Medium*

*5% sheep blood-GMM：Gut microbiota Medium+5% sterile defidrinated sheep blood*

*GMM：Gut microbiota Medium*

*5% sheep blood-R2A：R2A Medium+5% sterile defidrinated sheep blood*

*R2A：R2A Medium*

*5% sheep blood-SCH：SCH Medium+5% sterile defidrinated sheep blood*

*SCH：SCH Medium*

*5%sheep blood-**Spore：Spore Medium +5% sterile defidrinated sheep blood*

*Spore Medium*

*5% sheep blood- Columbia：Columbia Medium+5% sterile defidrinated sheep blood*

*Columbia Medium*

*Kana- Columbia：Columbia Medium+ Kanamycin*

*2216：DSMZ 2216 Medium*

*27：DSMZ 27 Medium*

*98：DSMZ 98 Medium*

*AM：AM medium*

*GYM：GYM Medium*

*NA：Nutrient Agar Medium*

1. ***Gut microbiota Medium***

| Component | Amount/L |
| --- | --- |
| Tryptone Peptone | 2 g |
| Yeast Extract | 1 g |
| D-glucose | 0.4 g |
| L-cysteine | 0.5 g |
| Cellobiose | 1 g |
| Maltose | 1 g |
| Fructose | 1 g |
| Meat Extract | 5 g |
| KH_2_PO_4_ | 100 mL |
| MgSO_4_·7H_2_O | 0.002 g |
| NaHCO_3_ | 0.4 g |
| NaCl_2_ | 0.08 g |
| CaCl_2_ | 1 mL |
| Vitamin K (menadine) | 1 mL |
| FeSO_4_ | 1 mL |
| Histidine Hematin Solution | 1 mL |
| Tween 80 | 2 mL |
| ATCC Vitamin Mix | 10 mL |
| ATCC Trace Mineral Mix | 10 mL |
| Acetic acid | 1.7 mL |
| Isovaleric acid | 0.1 mL |
| Propionnic acid | 2 mL |
| Butyric acid | 2 mL |
| Resazurin | 4 mL |
| Noble Agar | 12 g |
| pH | 7.2 |

2. *Spore Medium*

| Component | Amount/L |
| --- | --- |
| Yeast Extract | 1 g |
| Beef extract | 1 g |
| Tryptone Peptone | 2 g |
| Glucose | 10 g |
| FeSO_4_ | 0.001 g |
| Distilled water | 1000 mL |
| Noble Agar | 15 g |
| pH | 7.2 |

3. *DM Medium*

Solution A:

| Component | Amount/L |
| --- | --- |
| K_2_HPO_4_ | 0.5 g |
| NH_4_Cl | 1.0 g |
| Na_2_SO_4_ | 1.0 g |
| CaCl_2_·2H_2_O | 0.1 g |
| MgSO_4_·7H_2_O | 2.0 g |
| Na-DL-lactate | 2.0 g |
| Yeast extract | 1.0 g |
| Na-resazurin solution (0.1% w/v) | 0.5 mL |
| Distilled water | 980.0 mL |

Solution B:

| Component | Amount/L |
| --- | --- |
| FeSO_4_·7H_2_O | 0.5 g |
| Distilled water | 10 mL |

Solution C:

| Component | Amount/L |
| --- | --- |
| Na-thioglycolate | 0.1 g |
| Ascorbic acid | 0.1 g |
| Distilled water | 10 mL |

The solution B and solution C are added after the solution A has been boiled, adjusted pH to 7.8 with NaOH, Fixed capacity to 1000 mL, enter nitrogen to remove oxygen, Autoclave 15 min at 121℃.

4. ***R2A Medium***

| Component | Amount/L |
| --- | --- |
| Yeast extract | 0.5 g |
| Peptone | 0.5 g |
| Casein hydrolysate | 0.5 g |
| Glucose | 0.5 g |
| Starch, soluble | 0.5 g |
| K_2_HPO_4_ | 0.3 g |
| MgSO_4_ | 0.024 g |
| α-Ketopropionic acid sodium salt | 0.3 g |
| Noble Agar | 15.0 g |
| Distilled water | 1000 mL |
| pH | 7.2 ± 0.2 |

5. *Brain Heart Infusion Medium*

| Component | Amount/L |
| --- | --- |
| Dehydration Brain infusion powder | 12.50 g |
| Dehydration Beef Heart Infusio | 1.50 g |
| Proteose Peptone | 10.00 g |
| Glucose | 2.00 g |
| NaCl | 5.00 g |
| NaHPO_4_ | 2.50 g |
| Noble Agar | 15.00 g |
| pH | 7.4 |

6. *Columbia Medium*

| Component | Amount/L |
| --- | --- |
| Casein Tryptone | 10.0 g |
| Pepsin Hydrolytes | 5.0 g |
| Heart Pancreatin Hydrolytes | 3.0 g |
| Yeast Extract | 5.0 g |
| Corn starch | 1.0 g |
| NaCl | 5.0 g |
| Noble Agar | 15.0 g |
| Distilled water | 1000 mL |
| pH | 7.3 ± 0.2 |

7. *GYM Medium*

| Component | Amount/L |
| --- | --- |
| Glucose | 4.0 g |
| Yeast Extract | 4.0 g |
| Malt Extract | 10.0 g |
| CaCO_3_ | 2.0 g |
| Noble Agar | 12.0 g |
| Distilled water | 1000 mL |
| pH | 7.2 |

8. *AM medium*

| Component | Amount/L |
| --- | --- |
| KH_2_PO_4_ | 0.10 g |
| (NH_4_)_2_SO_4_ | 0.25 g |
| CaCl_2_·2H_2_O | 0.05 g |
| MgSO_4_·7H_2_O | 0.02 g |
| Trace elements | 1.0 mL |
| Yeast extract | 0.1 g |
| Na gluconate | 0.5 g |
| Distilled water | 1000 mL |
| Noble Agar | 15.0 g |
| pH | 5.0 - 5.5 |

9. *2216 Medium*

| Component | Amount/L |
| --- | --- |
| Peptone | 5.0 g |
| Yeast Extract | 1.0 g |
| Ferric Citrate | 0.1 g |
| NaCl | 19.45 g |
| MgCl_2_ | 8.8 g |
| Na_2_SO_4_ | 3.24 g |
| CaCl_2_ | 1.8 g |
| KCl | 0.55 g |
| NaHCO_3_ | 0.16 g |
| KBr | 0.08 g |
| SrCl_2_ | 34.0 mg |
| H_3_BO_3_ | 22.0 mg |
| Na_2_O·nSiO_2_ | 4.0 mg |
| NaF | 2.4 mg |
| NH_4_NO_3_ | 1.6 mg |
| Na_2_HPO_4_ | 8.0 mg |
| Noble Agar | 15.0 g |
| pH | 7.6 ± 0.2 |

10. *98 Medium*

| Component | Amount/L |
| --- | --- |
| Yeast extract | 1.0 g |
| Mannitol | 10.0 g |
| Noble Agar | 15.0 g |
| Soil extract | 200 mL |
| Distilled water | 800 mL |
| Noble Agar | 15.0 g |
| pH | 7.2 |

Soil extract:

| Component | Amount/L |
| --- | --- |
| Air-dried garden soil | 80.0 g |
| Na_2_CO_3_ | 0.2 g |
| Distilled water | 200 mL |

11. *27 Medium*

| Component | Amount/L |
| --- | --- |
| Yeast extract | 0.3 g |
| Na_2_-succinate | 1.0 g |
| (NH_4_)-acetate | 0.5 g |
| Fe(Ⅲ) citrate solution (0.1% in H_2_O) | 5 mL |
| KH_2_PO_4_ | 0.5 g |
| MgSO_4_·7H_2_O | 0.4 g |
| NaCl | 0.4 g |
| NH_4_Cl | 0.4 g |
| CaCl_2_·2H_2_O | 0.05 g |
| Vitamin B12 Solution (10 mg in 100 mL H_2_O) | 0.40 mL |
| Trace element solution SL-6 (See below) | 1.00 mL |
| L-Cysteine hydrochloride | 0.3 g |
| Resazurin (0.1%) | 0.5 mL |
| Distilled water | 1000 mL |
| Noble Agar | 15.0 g |
| pH | 6.8 |

12. *Nutrient Agar Medium*

| Component | Amount/L |
| --- | --- |
| Peptone | 5.0 g |
| beef extract | 30.0 g |
| NaCl | 5.0 g |
| Distilled water | 1000 mL |
| Noble Agar | 15.0 g |
| pH | 7.0 - 7.2 |

13. *SCH Medium*

| Component | Amount/L |
| --- | --- |
| Tryptone Peptone | 8.2 g |
| Peptone | 2.5 g |
| Peptone from soya | 1.0 g |
| Glucose | 5.8 g |
| Yeast extract | 5.0 g |
| NaCl | 1.7 g |
| NaHCO_3_ | 0.8 g |
| Cysteine-HCl·H_2_O | 0.4 g |
| Haemin | 0.01 g |
| Tris | 15.0 g |
| Distilled water | 1000 mL |
| Noble Agar | 15.0 g |
| pH | 7.2 |

14. *MPYG Medium*

| Component | Amount/L |
| --- | --- |
| Tryptone Peptone | 5.0 g |
| Peptone | 3.0 g |
| Peptone from soya | 2.0 g |
| Polypeptone | 1.0 g |
| Casein | 1.0 g |
| Yeast Extract | 10.0 g |
| Beef extract | 5.0 g |
| Glucose | 5.0 g |
| K_2_HPO_4_ | 2.0 g |
| Tween 80 | 0.5 mL |
| Maltose | 0.5 g |
| Cellobiose | 0.5 g |
| Starch, soluble | 0.5 g |
| Glycerol | 0.5 mL |
| Cysteine-HCl·H_2_O | 0.5 g |
| Na_2_S·9H_2_O | 0.25 g |
| Resazurin | 1.0 mg |
| Salt solution | 40.0 mL |
| Trace elements | 10.0 mL |
| Vitamin solution | 10.0 mL |
| Distilled water | 930 mL |
| Haemin solution | 10.0 mL |
| Vitamin K1 solution | 0.2 mL |
| Noble Agar | 15.0 g |
| pH | 7.0 |

The vitamin K1, Haemin solution, Trace elements, Vitamin solution and the cysteine are added after the medium has been boiled and cooled under CO_2_. Adjust pH to 7.0 using NaOH. Distribute under N_2_ and autoclave 15 min at 121℃.

Haemin solution:

Dissolve 50 mg Haemin in 1 mL 1 N NaOH; make up to 100 mL with distilled water. Store refrigerated.

Vitamin K1 solution:

Dissolve 0.1 mL of vitamin K1 in 20 mL 95% ethanol and filter sterilize. Store refrigerated in a brown bottle.
